# CREG1 restricts ALV-J replication via the mitochondrial dysfunction–driven activation of innate immunity and apoptosis

**DOI:** 10.1101/2025.08.18.670803

**Authors:** Qihong Zhang, Meihuizi Wang, Ming Pan, Junliang Xia, Tao Xu, Wen Luo, Xiquan Zhang

**Author notes:** Corresponding author. (X.Z.); (W.L.).

## Abstract

Despite the implementation of purification strategies to partially limit J subgroup avian leukosis virus (ALV-J) infection, the involvement of host factors in the underlying infection mechanism remains largely undefined. Here, we identify cellular repressor of E1A-stimulated genes 1 (CREG1) as a key regulator of mitochondrial function and a critical immune-related gene involved in ALV-J infection. The objective of this study was to explore the effects and underlying mechanisms of CREG1 in the context of ALV-J infection. Overexpression of *CREG1* upregulates the expression of type I interferon (I-IFN) and certain interferon-stimulated genes (ISGs), thereby suppressing viral replication. Mechanistically, overexpression of *CREG1* induces mitochondrial dysfunction, characterized by a decrease in mitochondrial membrane potential (Δψm), reduced adenosine triphosphate (ATP) production and respiratory chain activity, enhanced mitophagy, and increased release of mitochondrial DNA (mtDNA), which in turn triggers the activation of innate immune responses. Mitochondrial dysfunction further leads to the cytosolic release of cytochrome *c* and an increase in reactive oxygen species (ROS) levels, thereby triggering a robust apoptotic response. Moreover, the regulation of mitochondrial function by CREG1 depends on its interaction with the mitochondrial chaperone protein heat shock protein 1 (HSPD1), and their co-expression synergistically amplifies the antiviral response. In this study, we identify CREG1 as a potent antiviral gene and underscore the pivotal roles of mitochondria-mediated innate immunity and apoptosis during ALV-J infection.

**Author summary:** Despite efforts to limit J subgroup avian leukosis virus (ALV-J) infection, the role of host factors in the infection process is still not fully understood. In this study, we highlight the cellular repressor of E1A-stimulated genes 1 (CREG1) as a crucial regulator of mitochondrial function and immune responses during ALV-J infection. We show that CREG1 overexpression enhances type I interferon (I-IFN) and interferon-stimulated genes (ISGs) expression, inhibiting viral replication. This is achieved through mitochondrial dysfunction, including reduced mitochondrial membrane potential, lower ATP production, and increased mitophagy and mitochondrial DNA release, which activate innate immunity. CREG1-induced mitochondrial dysfunction also triggers apoptosis by releasing cytochrome c and increasing reactive oxygen species (ROS). Furthermore, CREG1 interacts with heat shock protein 1 (HSPD1) to amplify its antiviral effects. Our findings establish CREG1 as a key antiviral gene and emphasize the importance of mitochondrial-mediated immune responses and apoptosis in ALV-J infection.

## Introduction

Mitochondria, serving as the powerhouse of the cell, play a crucial role not only in cellular energy metabolism but also in viral infections and the innate immune response of the host[1]. Upon viral entry into the host cell, the virus often alters cellular metabolic pathways, particularly mitochondrial function. When mitochondria are damaged, they release various endogenous molecules, such as mitochondrial DNA (mtDNA), reactive oxygen species (ROS), and proteins, which have the capacity to activate the innate immune response[2, 3]. The release of mtDNA into the cytosol can be detected by cyclic GMP - AMP synthase (cGAS). Once recognized, cGAS catalyzes the production of cyclic GMP - AMP (cGAMP), a second - messenger molecule[4]. Subsequently, cGAMP activates the stimulator of interferon genes (STING)[5, 6]. This activation sets off a chain reaction that leads to the upregulation of both nuclear factor kappa - light - chain - enhancer of activated B cells (NF-κB) and proteins involved in the type I interferon (I-IFN) response[7]. In addition to innate immunity, host cells utilize apoptosis as an antiviral defense mechanism to eliminate infected or damaged cells[8]. When cells are subjected to stress or viral infection, mitochondria can initiate apoptosis via the mitochondrial pathway by modulating the B cell lymphoma 2 (BCL-2) family of proteins, leading to the release of pro-apoptotic factors such as cytochrome *c*[9, 10]. The increased permeability of the outer mitochondrial membrane facilitates the translocation of cytochrome *c* from the mitochondrial matrix into the cytosol, thereby activating the caspase family and triggering a cascade of signals that drive cell death[11, 12].

Avian leukosis virus (ALV), a retrovirus belonging to the family retroviridae, primarily infects avian species, especially chickens, and is closely associated with the onset of various immunosuppressive diseases and tumors[13]. Certain ALV subtypes, such as J subgroup ALV (ALV-J), are capable of inducing malignant transformation of host cells, leading to the development of lymphomas or other forms of tumors, thereby directly impacting poultry health and productivity[14]. In addition, the latent transmission characteristics of ALV mean that infected chickens do not exhibit obvious clinical symptoms initially, yet they can spread the virus throughout the entire farm through vertical transmission (from mother to offspring) or horizontal transmission (indirect transmission between chickens)[15]. Once clinical signs appear, it can cause significant economic losses to the poultry industry. Due to the variability of ALV and its immune evasion mechanisms, existing vaccines and antiviral drugs are often ineffective in preventing ALV-J infection[16, 17]. Therefore, gaining a deeper understanding of the interactions between the virus and the host immune system, identifying genes that inhibit ALV-J replication or enhance the host immune response, and improving overall immunity are key to addressing ALV infection.

In this study, we explored the role of cellular repressor of E1a-stimulating gene 1 (CREG1) during viral infection, based on its elevated expression in tissues and cells infected with ALV-J. CREG1 was initially discovered as a transcriptional repressor that antagonizes E1A oncoprotein 3-induced transcription and cell transformation[18]. Subsequent studies revealed that CREG1 is a secretory glycoprotein, widely expressed in adult tissues and cells, and is capable of promoting differentiation in human embryonal carcinoma cells[19]. Additionally, CREG1 is highly expressed in cardiac tissue and serves as a crucial myocardial protective factor[20]. Studies have demonstrated that CREG1 not only activates lysosomal autophagy to shield the heart from ischemia/reperfusion (MI/R) injury but also promotes the phenotypic transformation of myocardial fibroblasts after myocardial infarction by regulating the expression of cell division control protein 42 (CDC42)[21, 22]. Recent studies suggest that CREG1 enhances lysosome-associated membrane protein 2 (LAMP2) expression through an F-box protein 27 (FBXO27)-dependent pathway, thereby improving autophagy and mitigating the progression of diabetic cardiomyopathy[23]. In addition to its role in autophagy regulation, CREG1 has been found to localize to mitochondria, where it interacts with heat shock protein 1 (HSPD1) to regulate mitophagy, thereby maintaining skeletal muscle homeostasis[24]. Thus far, the biological role and underlying molecular mechanisms of CREG1 during ALV-J infection remain enigmatic.

Here, we demonstrate that the overexpression of CREG1 significantly inhibits the replication of ALV-J, primarily by enhancing innate immune responses. In vitro experiments demonstrated that overexpression of CREG1 impairs mitochondrial function, leading to the release of mtDNA. The released mtDNA subsequently activates the cGAS-STING signaling pathway, thereby enhancing I-IFN responses and the expression of interferon-stimulated genes (ISGs). Furthermore, CREG1 induces increased mitochondrial permeability, resulting in the release of cytochrome *c* into the cytoplasm, which activates the caspase cascade and executes apoptosis, serving as another critical factor in viral clearance. Finally, we identified an interaction between CREG1 and HSPD1, and their synergistic effect contributes to an enhanced antiviral response. Collectively, these findings provide new insights into the unique antiviral mechanism of CREG1 during ALV-J infection, advancing our understanding of potential strategies to enhance antiviral responses in poultry.

## Results

### CREG1 is a potential antiviral molecule against ALV-J infection

To investigate the key regulatory factors involved in the ALV-J infection process, we first performed bulk transcriptome sequencing on ALV-J-infected spleen tissues (Figures S1 and S2). In our previously published study, we confirmed that these samples were infected with ALV-J without cross-infection with other avian viruses[25]. These differentially expressed genes affect multiple signaling pathways, including immune regulation (e.g., Toll-like receptor signaling pathway and intestinal immune network for IgA production), cell signaling transduction (MAPK signaling pathway), and metabolic regulation (arachidonic acid metabolism, histidine metabolism, and nitrogen metabolism) (Figure 1A and 1B). These findings suggest that ALV-J infection may promote viral replication by affecting multiple key pathways, while also being associated with inflammation, immune suppression, or alterations in host physiological functions.

**Figure 1.**
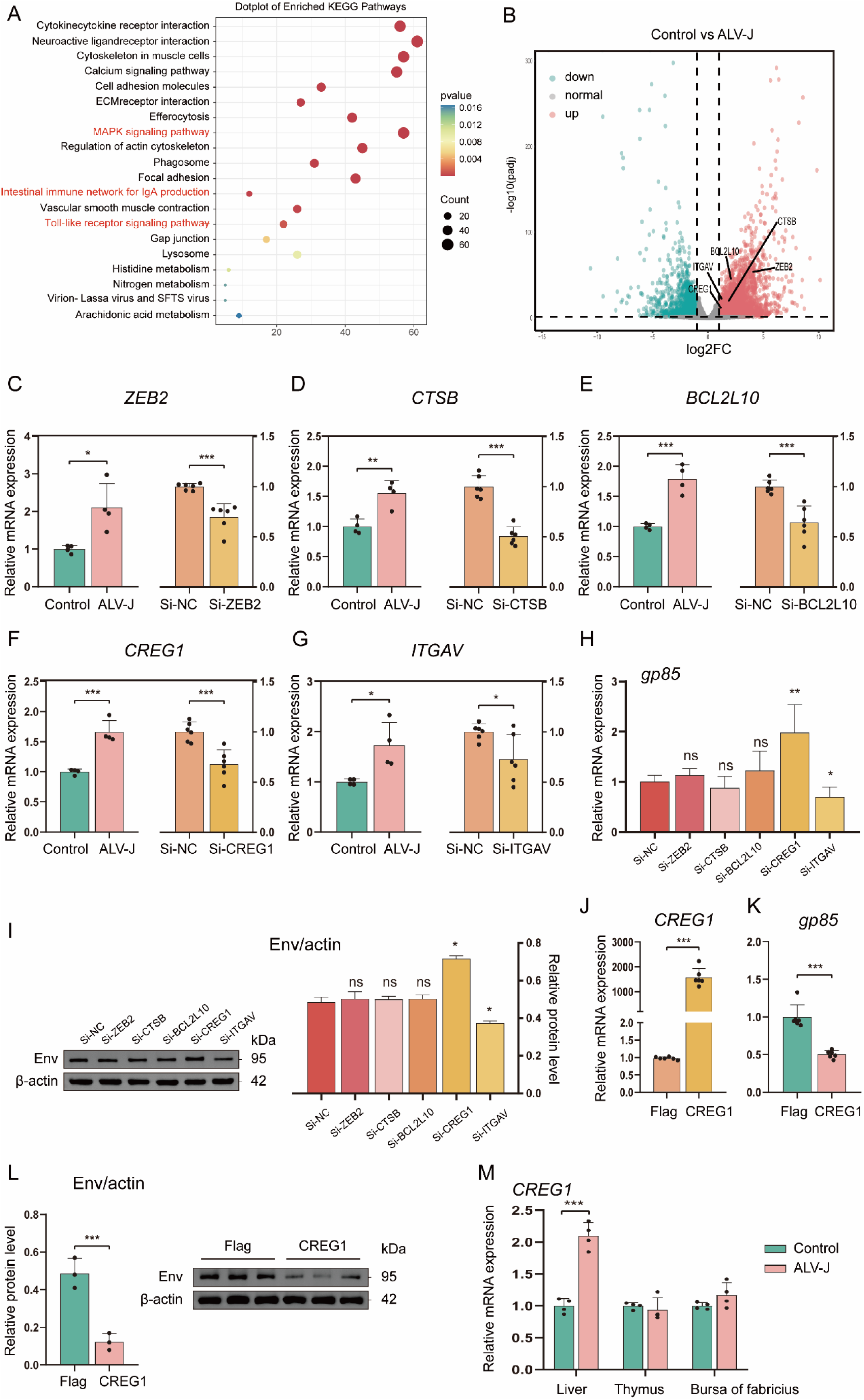
Overexpression of *CREG1* inhibits the replication of ALV-J. **(A).** Kyoto encyclopedia of genes and genomes (KEGG) enrichment analysis of differentially upregulated genes in RNA-seq data from ALV-J infected spleen and the control group. n = 4. **(B).** Volcano plot showing differentially expressed genes (DEGs) in the RNA-seq data from ALV-J-infected spleen tissues compared with the control group. DEGs were defined as those with |log₂ fold change| ≥ 1 and adjusted p-value ≤ 0.01. Red dots represent upregulated DEGs, while green dots represent downregulated DEGs. n = 4. **(C-G).** Relative mRNA expression levels of *ZEB2* (**C**), *CSTB* (**D**), *BCL2L10* (**E**), *CREG1* (**F**), and *ITGAV* (**G**) were sequentially detected by RT-qPCR. The left panel shows the expression levels of specific genes in the spleen tissue infected with ALV-J and the control group (n = 4). The right panel shows the expression levels of specific genes in DF-1 cells after 48 h of treatment with small interfering RNA (siRNA). n=6. **(H).** After siRNA treatment of specific genes in DF-1 cells, the cells were infected with ALV-J. The expression level of the viral protein gp85 was measured by RT-qPCR 48 hours post-infection (hpi). n = 6. **(I).** After siRNA treatment of specific genes in DF-1 cells, the cells were infected with ALV-J. At 48 hpi, the expression levels of the viral envelope protein env and β-actin were detected by western blotting. n = 3. The left panel shows the immunoblot, and the right panel shows the relative expression levels of env (env/actin). **(J).** After transfection of Flag-CREG1 or control (Flag-pCDNA3.1 without insert was used as a negative control.) into DF-1 cells, the expression level of *CREG1* was measured by RT-qPCR at 48 hpi. n = 6. **(K).** DF-1 cells were transfected with Flag-CREG1 or control, followed by ALV-J infection. The expression level of the viral protein gp85 was then assessed by RT-qPCR at 48 hpi. n = 6. **(L).** Flag-CREG1 or control was transfected into DF-1 cells, followed by ALV-J infection. At 48 hpi, the expression levels of the viral envelope protein env and β-actin were analyzed by western blotting. The left panel shows the immunoblot, and the right panel shows the statistical analysis of relative env expression (env/actin). n = 3. **(M).** RT-qPCR was used to measure the relative expression levels of *CREG1* in the liver, thymus, and bursa of Fabricius following ALV-J infection or in the control group. n = 4. For **C-M**, the data are presented as mean ±SD. Statistical significance was determined using unpaired Student’s t-test, with ns indicating no significant difference, *p < 0.05, **p < 0.01, and ***p < 0.001.

Next, we investigated the impact of several upregulated genes, including *ZEB2*, *BCL2L10*, *ITGAV*, *CTSB*, and *CREG1*, on viral replication. Notably, these genes were also highly expressed in certain immune cell subsets in single-cell sequencing data of ALV-infected peripheral blood mononuclear cells[26]. The expression of these genes was induced in the ALV-J-infected spleen, as confirmed by RT-qPCR (Figure 1C-1G). To verify the effect of these genes on viral replication efficiency, we performed gene knockdown using small interfering RNA (siRNA) in the chicken fibroblast cell line (DF-1), followed by infection with ALV-J for 48 h. The knockdown efficiency was detected by RT-qPCR (Figure 1C-1G). We found that depletion of CREG1 significantly enhanced viral replication, while depletion of ITGAV notably suppressed viral replication. No significant effect on viral replication was observed for the other genes (Figure 1H and 1I). To further confirm the roles of CREG1 and ITGAV in viral replication, we constructed expression plasmids for Flag-CREG1 and Flag-ITGAV (Figure 1J and Figure S3A). The results demonstrated that *CREG1* overexpression (OE-CREG1) markedly inhibited viral replication, while *ITGAV* overexpression (OE-ITGAV) markedly promoted viral replication (Figure 1K-1L and Figure S3B-S3D). Thus, these results suggest that CREG1 may be a potential antiviral molecule in the course of ALV-J infection.

Furthermore, by analyzing the expression of CREG1 in chicken tissues, we found that the mRNA level of *CREG1* was also significantly higher in the liver tissues infected with ALV-J compared to the healthy group (Figure 1M). Similarly, we obtained the upregulated expression pattern of *CREG1* in DF-1 cells, primary chicken fibroblasts (CEF), MSB-1 cells, or chicken B lymphocyte (DT40) cells under ALV-J infection through RT-qPCR analysis (Figure S4A). These results suggest that ALV-J infection induces the upregulation of *CREG1* expression.

### CREG1 initiates innate immune responses

To further explore the role of CREG1 in antiviral immune responses, we performed quantitative proteomics analyses on *CREG1*-overexpressing ALV-J-infected DF-1 cells. Surprisingly, CREG1 induced the expression of a significant number of ISGs, including IFIT5, MX1, ZNFX1, CCL4, ISG12(2), OASL and others (Figure 2A and Figure S5A-S5B). Strikingly, in this proteomics data, we found that the gene encoding the ALV-J envelope glycoprotein, env, appeared in the downregulated group, further confirming the inhibitory effect of CREG1 on the virus (Figure 2A and Figure S5A-S5B). We confirmed the induction of certain ISGs in *CREG1*-overexpressing ALV-J-infected cells using RT-qPCR (Figure 2B and 2C). Upregulation of ISG expression suggests that CREG1 may have activated the interferon pathway. In fact, in the results of the quantitative protein analysis, we also found that interferon induced with helicase C domain 1 (IFIH1, also known as MDA5) and STING1 were upregulated (Figure 2A). Activation of MDA5 or STING promotes the production of I-IFN, further activating the antiviral immune response[27]. To verify this hypothesis, we examined the activation of the interferon pathway after transfecting Flag-CREG1 and subsequently infecting with ALV-J. The results indicated that in DF-1 and HD11 cells (chicken macrophage cell line), OE-CREG1 induced the activation of IFN-β through the cGAS-STING pathway (Figure 2D-2F), accompanied by key downstream signaling events, including phosphorylation of Janus kinase (JAK) and signal transducer and activator of transcription 1 (STAT1) (Figure 2G-2H and Figure S5C). Although the overexpression of *CREG1* did not cause differential expression of *MDA5* (Figure S5D). Interestingly, when we stimulated the cells with I-IFN, the expression of *CREG1* did not significantly increase, suggesting that CREG1 is not a classic ISG (Figure S5E). Moreover, in the single-cell omics data, CREG1 is highly expressed in the Th1-like cell subpopulation[26]. We speculate that CREG1 might also activate the antiviral immune response of T cells. To verify this, we transfected Flag-CREG1 into the chicken T lymphocyte cell line MSB-1, followed by ALV-J infection, and subsequently evaluated the secretion of IFN-γ, interleukin-10 (IL-10), and IL-4. Unexpectedly, the results showed that CREG1 did not effectively activate T cells (Figure S5F). Meanwhile, the deletion of *CREG1* significantly weakened the production of IFN-β and the expression of ISGs (Figure S5G and S5H). Overall, these data suggest that CREG1 initiates the innate immune responses.

**Figure 2.**
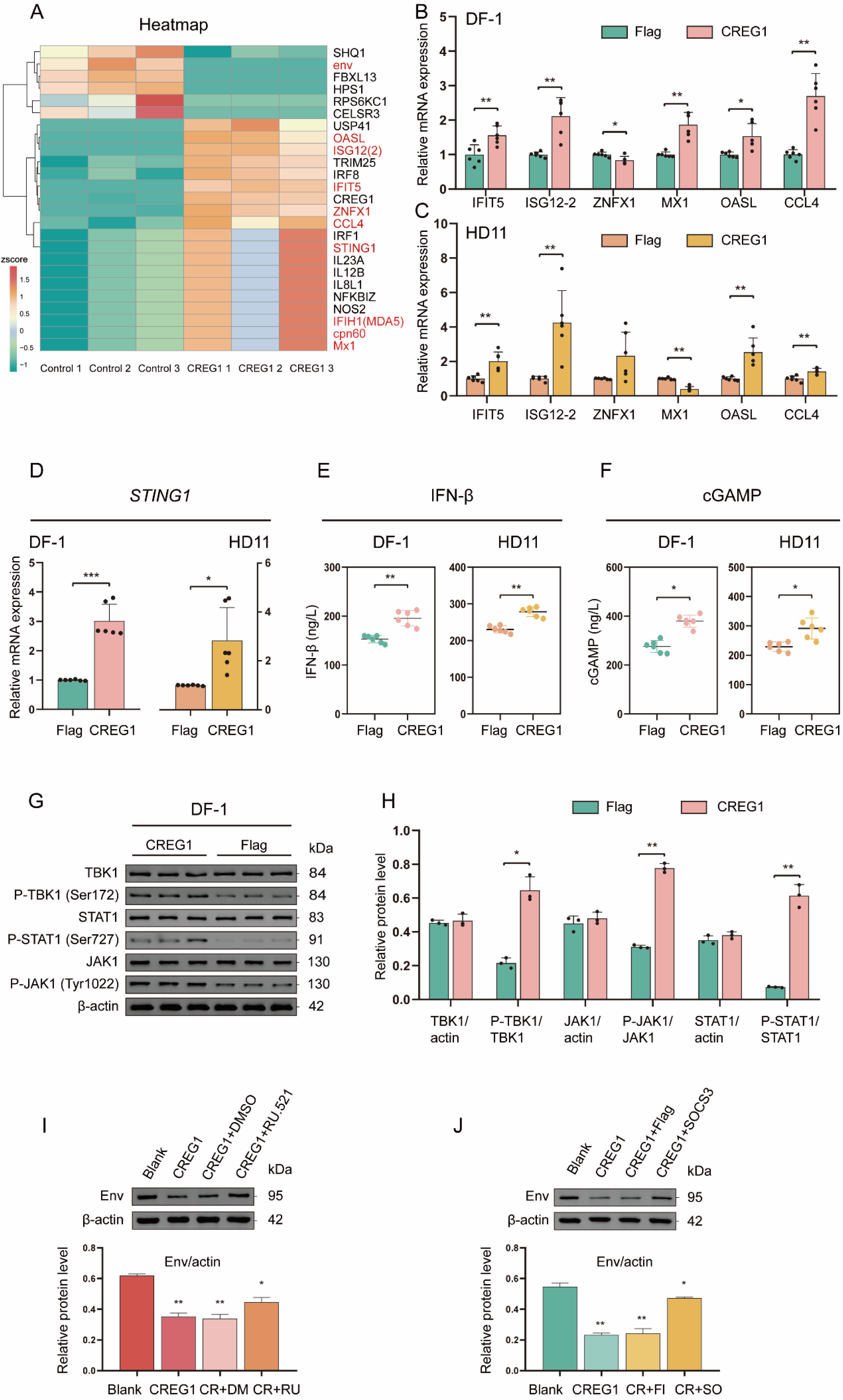
CREG1 triggers innate immune response activation. **(A).** Overexpression of *CREG1* or control in DF-1 cells infected with ALV-J, with proteomic results at 48 hpi. The heatmap shows the results of several key differential proteins, including ISGs and the viral protein env. Z-score normalized log2(FPKM) values, n = 3. **(B-C).** Overexpression of *CREG1* or control in DF-1 (**B**) or HD11 (**C**) cells infected with ALV-J, with RT-qPCR analysis of ISGs mRNA expression at 48 hpi, including *IFIT5*, *ISG12-2*, *ZNFX1*, *MX1*, *OASL*, and *CCL4*. n = 6. **(D).** Overexpression of *CREG1* or control in DF-1 (left) or HD11 (right) cells followed by ALV-J infection, with *STING1* mRNA expression levels measured by RT-qPCR at 48 hpi, n = 6. **(E-F).** *CREG1* or control overexpression in DF-1 (left) or HD11 (right) cells, followed by ALV-J infection, with cell supernatants harvested at 48 hpi for IFN-β and cGAMP quantification by enzyme-linked immunosorbent assay (ELISA), n = 6. **(G-H).** *CREG1* or control overexpression in DF-1 followed by ALV-J infection, with western blot analysis at 48 hpi to assess the activation of the I-IFN signaling pathway, including TBK1, P-TBK1 (Ser172), STAT1, P-STAT1 (Ser727), JAK, P-JAK (Tyr1022), and actin. The left panel shows the immunoblotting results **(G)**, while the right panel presents the densitometric analysis of relative protein expression **(H)**. n = 3. **(I).** DF-1 cells were treated with blank, *CREG1* overexpression alone, *CREG1* + DMSO, or *CREG1* + RU.521 (5 μM RU.521 for 2 h), followed by ALV-J infection. At 48 hpi, western blot was used to measure the viral protein env and actin. The upper panel shows the immunoblotting results, while the lower panel presents the densitometric analysis of relative protein expression. n = 3. **(J).** DF-1 cells were subjected to blank treatment, *CREG1* overexpression alone, *CREG1* + FLAG, or *CREG1* + *SOCS3* co-transfection, followed by ALV-J infection. At 48 hpi, viral envelope protein (env) and actin were assessed using western blot. The upper panel shows the immunoblotting results, while the lower panel presents the densitometric analysis of relative protein expression. n = 3. For **B-F and H-J**, the values are shown as the mean ± SD. *p < 0.05, **p < 0.01, ***p < 0.001 by unpaired Student’s t test.

To further investigate whether the antiviral effect of CREG1 depends on the innate immune responses, we pretreated OE-CREG1 cells with the cGAS inhibitor, RU.521, followed by virus infection. Interestingly, when the type Ⅰ-IFN response is inhibited, the suppressive effect of CREG1 on viral replication was partially diminished (Figure 2I). Meanwhile, since suppressor of cytokine signaling 3 (SOCS3) negatively regulates the JAK-STAT pathway[28], we found that co-transfection of *SOCS3* and *CREG1* resulted in a partial attenuation of the antiviral activity of CREG1 (Figure 2J). Therefore, we conclude that the antiviral effect of CREG1 is partially dependent on the innate immune response induced by the cGAS-STING-TBK1 pathway.

### CREG1 induces mitochondrial dysfunction

Next, we begin to investigate the reasons for CREG1 activation of the innate immune response. Mitochondria are not only the energy metabolism factories of the cell, but also, when mitochondrial dysfunction occurs, they release mtDNA, which activate the STING pathway, promote the production of I-IFN, and subsequently initiate innate immune responses and regulate the activity of immune cells[29, 30]. Meanwhile, previous reports have identified that CREG1 localizes to mitochondria and regulates the motility of skeletal muscle through mitochondrial autophagy[24]. Therefore, we hypothesize that CREG1 may trigger innate immune pathways by affecting mitochondrial function. To this end, we transfected Flag-CREG1 into DF-1 cells and examined its subcellular localization by immunofluorescence, with or without ALV-J infection. However, we found that CREG1 was not localized to mitochondria in chicken cells. Instead, the exogenously introduced CREG1 was mainly distributed in the cytoplasm and nucleus (Figure S6A). We then investigated whether the overexpression of *CREG1* affects mitochondrial biological function under viral infection conditions. Transmission electron microscopy (TEM) analysis showed that the number of abnormal mitochondria was significantly increased in OE-CREG1 cells (Figure 3A-3C). Compared to the control group, OE-CREG1 cells exhibited mitochondrial damage, such as cristae shortening and disintegration, reduced matrix electron density, and mitophagy (Figure 3A-3C). Mitochondrial membrane potential (TMRE and JC-1 staining), mitochondrial ROS (MitoSOX) were also assessed in OE-CREG1 DF-1 cells by flow cytometry (Figure 3D-3I). The results showed that CREG1 induced a significant decrease in mitochondrial membrane potential (Figure 3G and 3I). Furthermore, OE-CREG1 cells exhibited significantly lower mitochondrial superoxide levels, as detected by MitoSOX staining (Figure 3H). To determine whether mitochondrial dysfunction had occurred, we evaluated key mitochondrial functions such as ATP synthesis and respiratory chain activity. As expected, a reduction in ATP synthesis capacity and respiratory chain activity was observed in OE-CREG1 DF-1 cells compared with the control group (Figure 3J-3L).

**Figure 3.**
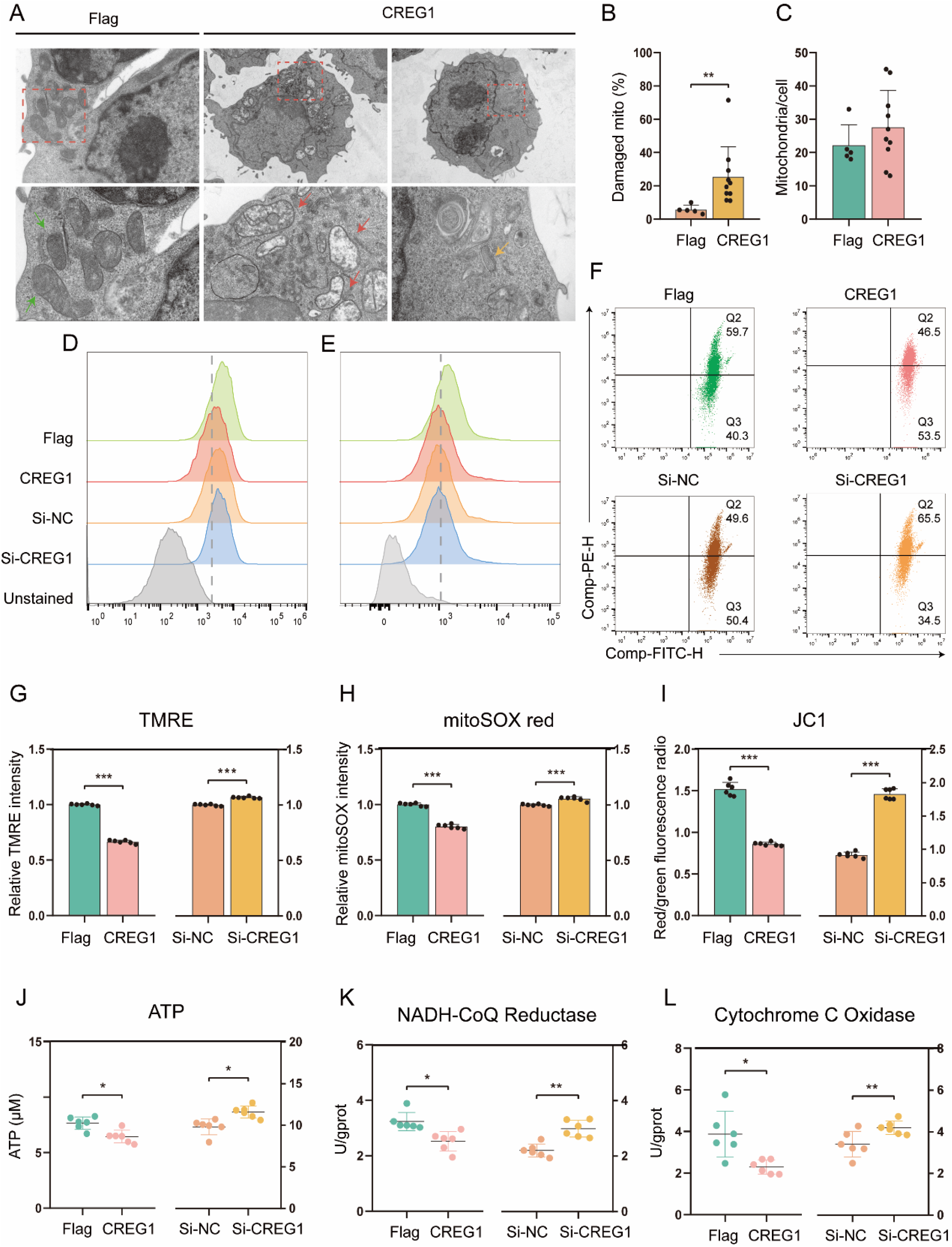
CREG1 disrupts mitochondrial integrity and function. **(A-C).** Transmission electron microscopy (TEM) images of HD11 cells transfected with Flag-CREG1 or control and infected with ALV-J for 48 hpi (**A**). The lower panels show magnified views of the corresponding upper panels. Green arrows indicate morphologically normal mitochondria; red arrows indicate swollen mitochondria or those with cristae loss; yellow arrows indicate mitophagy. Scale bars: 1.0 μm or 5.0 μm (upper panels), 500 nm or 1.0 μm (lower panels). Quantification of the number of mitochondria (**B**) and the percentage of damaged mitochondria (**C**). **(D-F).** Flow cytometry histograms or dot plots of TMRE (**D**), MitoSOX (**E**), and JC-1 (**F**) staining in *CREG1*-overexpressing or knockdown cells and corresponding control groups at 48 h post ALV-J infection. n = 6. **(G-H).** Quantification of relative fluorescence intensity of TMRE (**G**) and MitoSOX staining (**H**). **(I).** The bar graph represents the ratio of red fluorescence to green fluorescence. **(J-L).** After 48 h of ALV-J infection, ATP (**J**), NADH-CoQ Reductase (**K**), and Cytochrome C Oxidase (**L**) levels were measured in *CREG1*-overexpressing or knockdown cells and the control group, respectively. n = 6. For **B, C**, and **G-L**, the values are shown as the mean ± SD. *p < 0.05, **p < 0.01, ***p < 0.001 by unpaired Student’s t test.

Mitochondrial dysfunction may trigger mitophagy to remove damaged mitochondria. We analyzed the status of mitophagy under in OE-CREG1 or Si-CREG1 DF-1 cells. We performed subcellular fractionation of mitochondria and cytoplasm and found that ectopic expression of *CREG1* led to an upregulation of Parkin (RBR E3 ubiquitin protein ligase) in the mitochondrial fraction, accompanied by a decrease in the outer membrane protein TOMM20 (Figure 4A and 4B). In contrast, Parkin expression in the cytoplasmic fraction was reduced (Figure 4A and 4B). Furthermore, analysis of the autophagy-related proteins LC3 and p62 in total cellular lysates revealed that ectopic expression of *CREG1* increased LC3-II levels and decreased p62 levels, whereas siRNA-mediated knockdown of *CREG1* produced the opposite effects (Figure 4E). These results suggest that CREG1 may induce mitophagy. To confirm the findings, we further assessed the mitophagy flux in OE-CREG1 cells. Colocalization analysis of GFP-LC3 with MitoTracker Deep Red (MTDR) showed that CREG1 promotes the delivery of LC3B to mitochondria (Figure S6B). Taken together, these results indicate that the presence of CREG1 promotes mitochondrial dysfunction and subsequently induces mitophagy.

**Figure 4.**
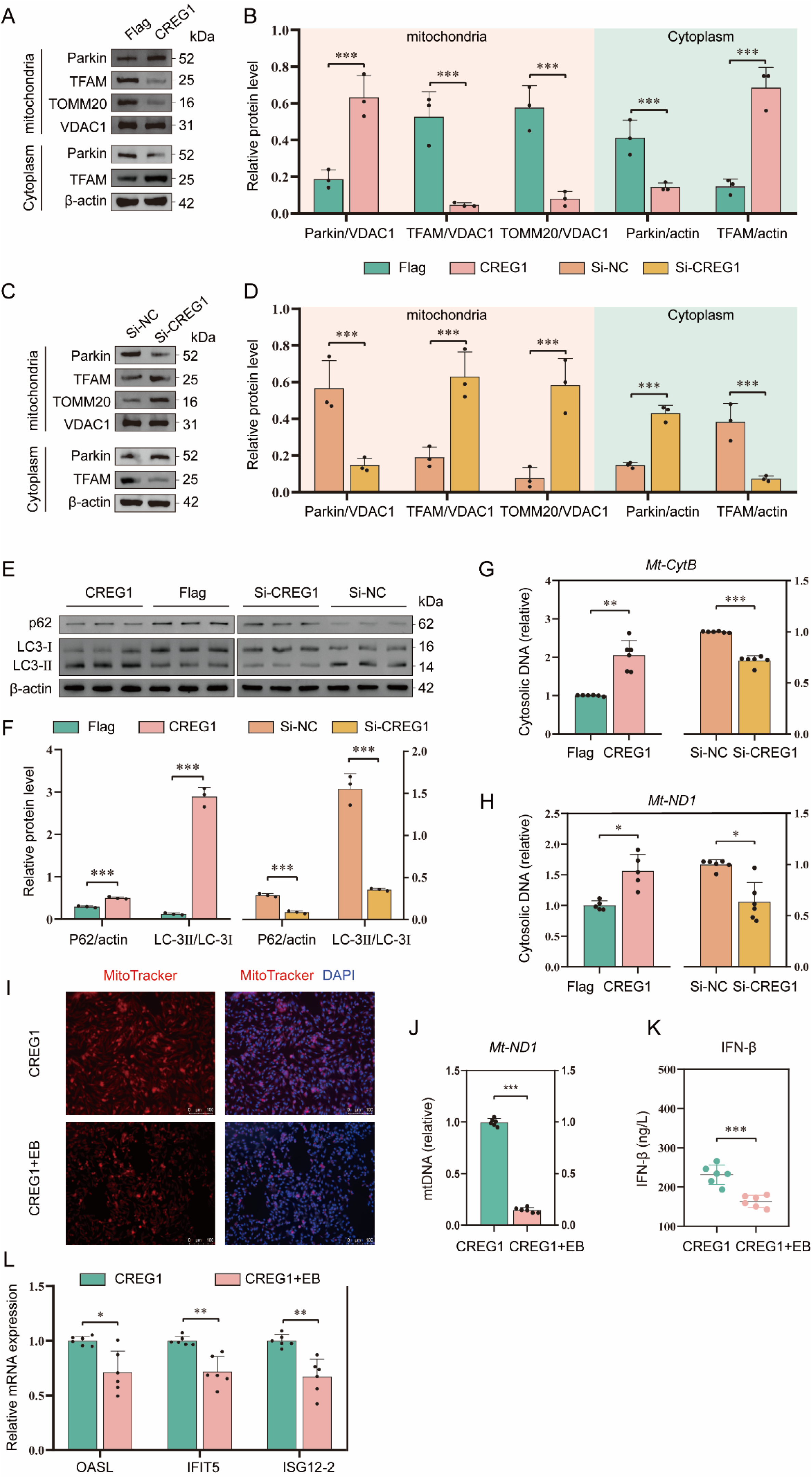
CREG1 induces mtDNA release into the cytosol and promotes mitophagy. **(A-D).** DF-1 cells with *CREG1* overexpression (**A-B**) or knockdown (**C-D**) were infected with ALV-J. At 48 hpi, cells were harvested and subjected to mitochondrial and cytosolic fractionation. The expression of Parkin, TFAM, TOMM20, and VDAC1 in the mitochondrial fraction, as well as Parkin and TFAM in the cytosolic fraction, was analyzed by western blotting. n = 3. **(E-F).** The expression levels of P62 and LC3-I/II in total cellular proteins from *CREG1*-overexpressing or *CREG1*-knockdown cells were analyzed by western blotting. Immunoblot image (**E**), statistical grayscale plot (**F**). n = 3. **(G-H).** Cells overexpressing or silencing *CREG1* were infected with ALV-J and subjected to subcellular fractionation. The cytosolic fraction was collected for DNA extraction, and the cytosolic mtDNA levels were quantified by RT-qPCR. n = 6. **(I-J).** *CREG1*-overexpressing cells were left untreated (UNT) or treated with ethidium bromide (EB) for 5 d. Mitochondria and nuclei were labeled with MitoTracker Red and DAPI, respectively (**I**). Scale bars, 100 µm. RT-qPCR detection of total cellular mtDNA expression levels (**J**). n = 6. **(K).** The level of IFN-β in the supernatant of untreated or EB-treated *CREG1*-overexpressing cells was measured by ELISA. n = 6. **(L).** The expression levels of *OASL*, *IFIT5*, and *ISG12-2* in untreated or EB-treated *CREG1*-overexpressing cells were quantified by RT-qPCR. n = 6. For **B, D, F, G, H, J, K, L,** the values are shown as the mean ± SD. *p < 0.05, **p < 0.01, ***p < 0.001 by unpaired Student’s t test.

Subsequently, we sought to reassess mitochondrial function by knocking down *CREG1* using siRNA. In contrast to the results obtained from OE-CREG1 cells, the absence of *CREG1* led to enhanced ATP production and respiratory chain activity, accompanied by a reduction in mitophagy (Fig 3J-3L and 4C-4E). Notably, upon revisiting the proteomics data, we found that the antiviral response of CREG1 was associated with certain mitochondrial proteins, such as heat shock protein family D (Hsp60) member 1 (HSPD1, also known as cpn60), a mitochondrial chaperonin involved in protein folding. HSPD1 is upregulated in response to cellular stress and may serve as an early indicator of mitochondrial damage[31]. Overall, these data indicate that during antiviral responses, CREG1 induces mitochondrial morphological damage, disrupts electron transport, ATP production, and membrane potential, and promotes mitophagy.

### CREG1 induces the release of mtDNA into the cytosol

Mitochondrial dysfunction triggers innate immune signaling by releasing mtDNA into the cytosol. To investigate whether CREG1 affects the release of mitochondrial mtDNA, we analyzed the accumulation of mtDNA in the cytosol of OE-CREG1 cells. Our results showed that ectopic *CREG1* expression resulted in marked cytosolic accumulation of mitochondrial genomic DNA (Figure 4G and 4H). Concurrently, an accumulation of mitochondrial transcription factor A (TFAM) was also detected in the cytosol (Figure 4A and 4B). However, no significant increase in mtRNA levels was observed in OE-CREG1 cells relative to the control group (Figure S6C-S6D).

Next, to determine whether the innate immune response observed in OE-CREG1 cells depends on mtDNA, we employed ethidium bromide (EtBr) to deplete mitochondrial nucleic acids. The results showed that EtBr effectively removed mitochondrial nucleic acids from the cells and markedly suppressed the expression of I-IFN and ISGs in *CREG1*-overexpressing cells (Figure 4I-4L). These results suggest that mtDNA in the cytosol may be key mediators of the CREG1-induced innate immune response.

### The pro-apoptotic effect of CREG1 is essential for its antiviral activity

Mitochondria act as a central regulator of multiple cell death pathways. Through the regulation of cytochrome *c* release, membrane potential, ROS generation, and mitochondrial permeability transition pore (mPTP) opening, they critically determine cell fate[32]. We propose that CREG1-induced mitochondrial dysfunction may further trigger programmed cell death during antiviral responses. To this end, we measured intracellular levels of ROS and NO, as well as the release of cytochrome *c* into the cytosol. The results showed that ectopic expression of *CREG1* enhanced ROS production (indicated by increased DCFDA fluorescence) and NO generation, and promoted the accumulation of cytosolic cytochrome *c* (Figure 5A-5D). In contrast, knockdown of *CREG1* reversed these effects, suggesting that CREG1 may induce cell death (Figure 5A-5D). Our initial hypothesis was that CREG1 induced disease-associated inflammatory pyroptosis. Unexpectedly, we did not observe an upregulation of IL-1β and TNF-α in the supernatant of OE-CREG1 cells, suggesting that inflammatory pyroptosis did not occur (Figure S7A and S7B). Subsequently, apoptosis was assessed by flow cytometry. The results demonstrated that ectopic expression of *CREG1* significantly enhanced cell apoptosis under viral infection (Figure 5E and 5F). Consistently, analysis of the apoptosis-related proteins BCL-2, BAX, and caspase-3 further confirmed that CREG1 promotes apoptosis during antiviral responses (Figure 5G). Conversely, the absence of *CREG1* significantly attenuated apoptosis (Figure S7C-S7E). Taken together, these findings suggest that CREG1 plays a pro-apoptotic role.

**Figure 5.**
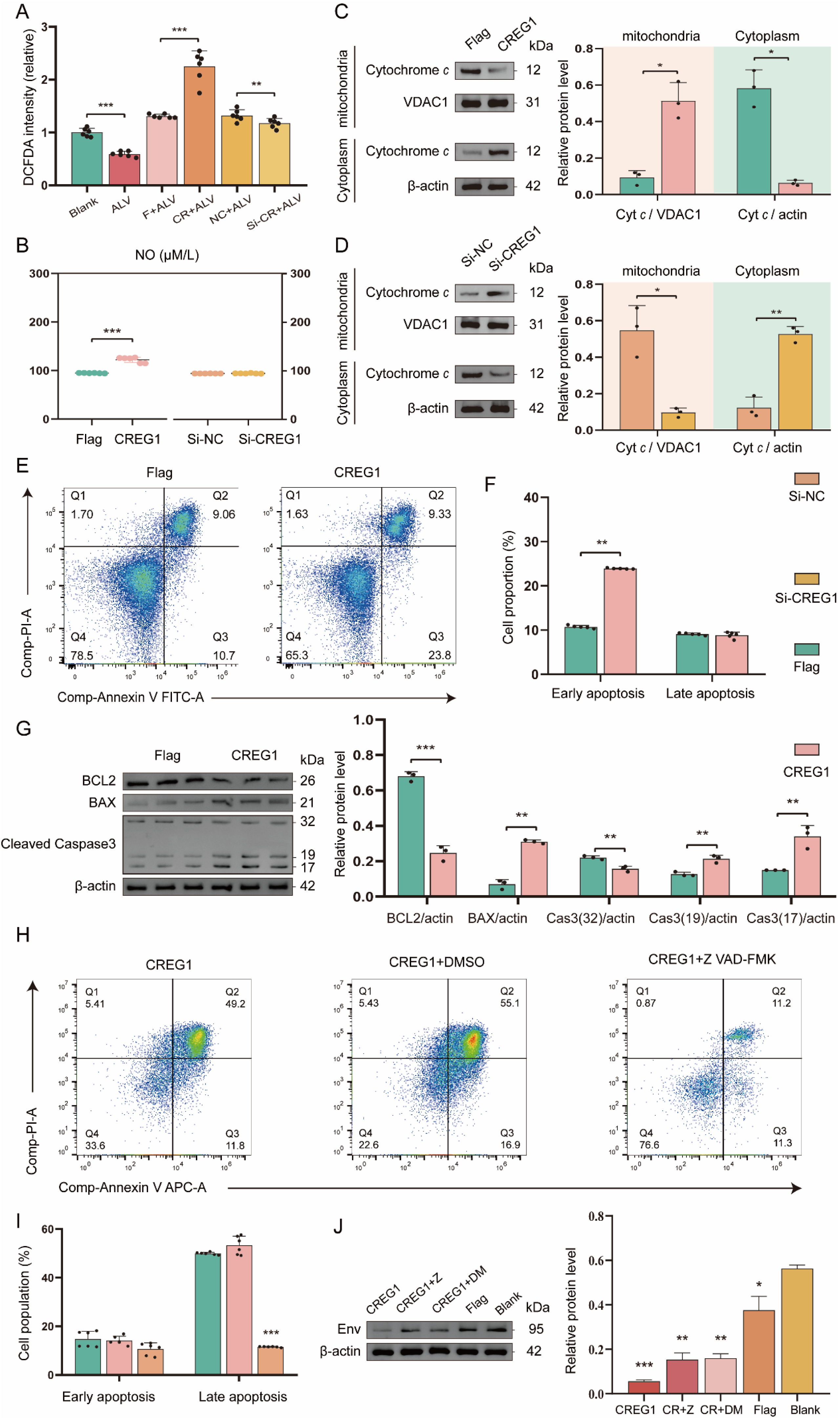
CREG1-mediated pro-apoptotic activity contributes to its antiviral function. **(A-B).** DF-1 cells with *CREG1* overexpression or knockdown were infected with ALV-J, and at 48 hpi, cells were collected for DCFDA (**A**) and NaNO_2_ (**B**) fluorescence measurement using a microplate reader. n = 6. **(C-D).** Western blot was performed to detect cytochrome *c* levels in mitochondrial and cytosolic fractions of *CREG1*-overexpressing (**C**) or -knockdown cells (**D**) at 48 hpi with ALV-J. n = 3. **(E-F).** Apoptosis was analyzed by Annexin V/PI staining and flow cytometry in *CREG1*-overexpressing and control cells infected with ALV-J. (**E**) Flow cytometry dot plots illustrating apoptotic cell distribution. (**F**) Statistical analysis of early and late apoptosis. n = 5. **(G).** Western blot analysis of apoptosis-related proteins (BCL2, BAX, and cleaved caspase-3) in *CREG1*-overexpressing and control cells upon ALV-J infection. The left panel shows representative immunoblots, and the right panel presents the quantification of relative protein expression levels. n = 3. **(H-I).** Cells were transfected to overexpress *CREG1* for 24 h and then pretreated with Z-VAD-FMK (50 μM) or DMSO for 1 h prior to ALV-J infection. Apoptosis was analyzed by flow cytometry 24 hpi. (**H**) Representative flow cytometry dot plots. **(I).** Quantification of early and late apoptotic cells. n = 6. **(J).** *CREG1*-overexpressing cells were pretreated with Z-VAD-FMK (50 μM) or DMSO for 1 h prior to ALV-J infection following 24 h of transfection. Viral envelope protein env expression was analyzed by Western blot at 24 hpi. The left panel shows representative immunoblots, and the right panel presents the quantification of relative protein expression levels. n = 3. For **A-D, F, G, I, J,** the values are shown as the mean ± SD. *p < 0.05, **p < 0.01, ***p < 0.001 by unpaired Student’s t test.

Subsequently, to investigate whether the antiviral activity of CREG1 is linked to its pro-apoptotic effect, we treated OE-CREG1 cells with an apoptosis inhibitor, Z-VAD-FMK, prior to ALV-J infection. Treatment with Z-VAD-FMK significantly inhibited CREG1-induced cell apoptosis but had no effect on the activation of I-IFN (Figure 5H-5I and Figure S7F). Next, we investigated the impact of CREG1 on viral replication when apoptosis was suppressed. The results demonstrated that the antiviral activity of CREG1 was markedly compromised under conditions of apoptosis inhibition (Figure 5J).

### CREG1 interacts with HSPD1, and its regulatory effect on mitochondrial function is contingent upon HSPD1

To further investigate the mechanism by which CREG1 regulates mitochondrial dysfunction, we employed co-immunoprecipitation and mass spectrometry to screen for proteins interacting with CREG1 under viral infection. Notably, mass spectrometry analysis revealed that many of the potential CREG1-interacting proteins are involved in mitochondrial function, including molecular chaperones (e.g., HSPD1, HSPA9), respiratory chain-related proteins (e.g., SDHA, UQCRC2), transporter-related proteins (e.g., SLC25A3, SLC25A4), and structural stability proteins (e.g., PHB, PHB2) (Supplementary Table 4). Previous reports demonstrated an interaction between CREG1 and HSPD1 in mouse skeletal muscle[24]. In our previous proteomic analysis, HSPD1 was identified as upregulated. Consistently, western blot analysis confirmed that overexpression of *CREG1* led to an increase in HSPD1 expression, whereas knockdown of *CREG1* resulted in its downregulation (Figure S8A and S8B). These findings imply a potential interaction between CREG1 and HSPD1. To this end, we transfected Flag-CREG1 alone or co-transfected it with HA-HSPD1 into DF-1 cells to determine whether an interaction between the two proteins occurs in the presence or absence of ALV-J infection. As expected, co-immunoprecipitation assays confirmed that CREG1 interacts with either endogenous HSPD1 or exogenously expressed HA-HSPD1 under both infected and uninfected conditions (Figure 6A and 6B). Therefore, these findings indicate that CREG1 interacts with HSPD1.

**Figure 6.**
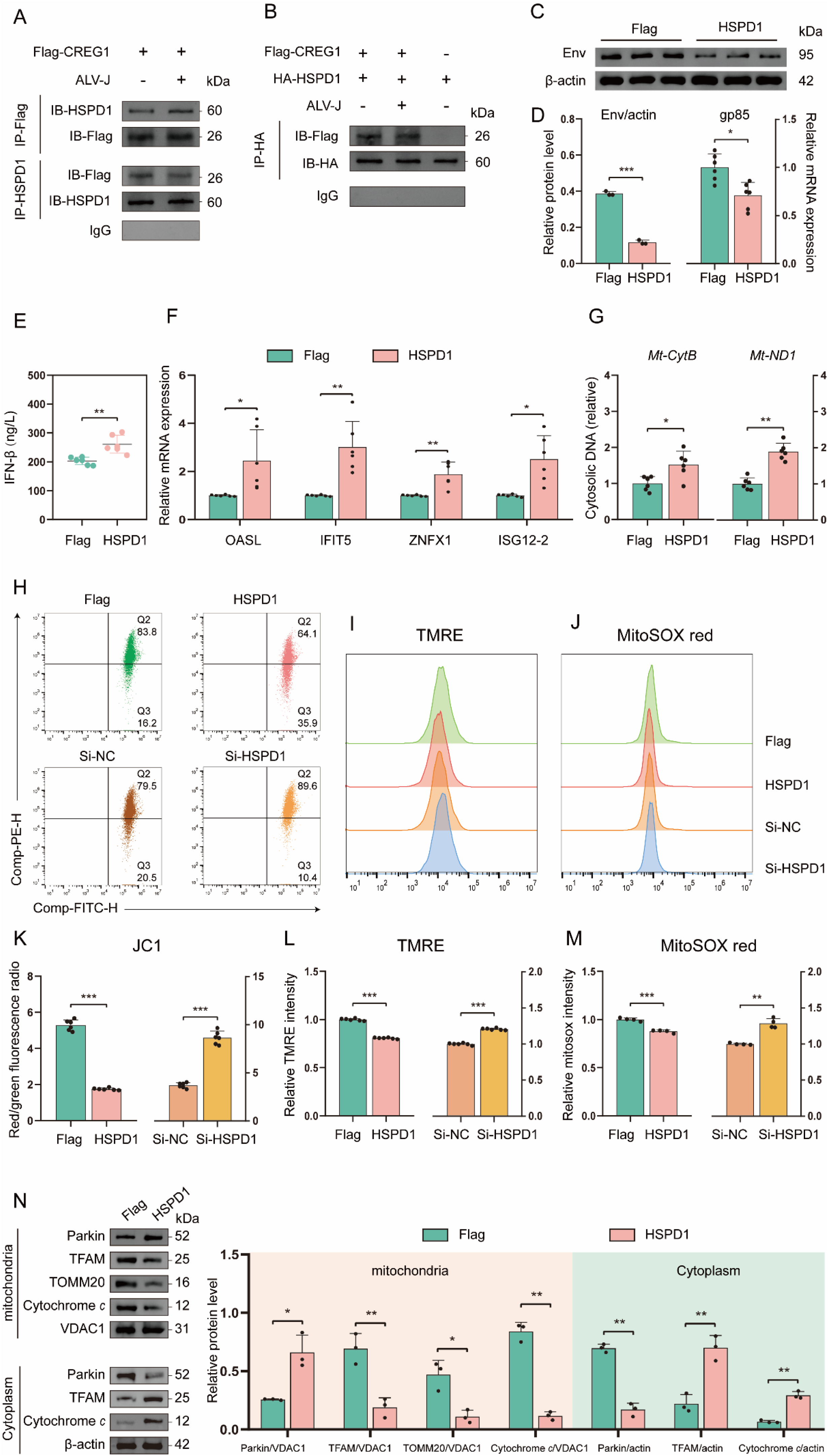
CREG1 interacts with HSPD1, and HSPD1 also exhibits antiviral activity. **(A-B).** Co-IP analysis of CREG1–HSPD1 interaction with or without ALV-J infection. **(C-D).** ALV-J expression was detected by western blot (**C**) or RT-qPCR (**D**) at 48 hpi following *HSPD1* overexpression. n = 3 (**C**) or n = 6 (**D**). **(E-G).** ELISA detection of IFN-β in the supernatant, and RT-qPCR analysis of ISGs (*OASL*, *IFIT5*, *ZNFX1*, *ISG12-2*) expression and cytosolic mtDNA levels in *HSPD1*-overexpressing cells at 48 hpi with ALV-J. n = 6. **(H-M).** In ALV-J-infected cells with *HSPD1* overexpression or knockdown, the fluorescence intensities of JC1 (**H**), TMRE (**I**), and MitoSOX (**J**) were measured using a microplate reader at 48 hpi. The ratio of red fluorescence to green fluorescence (**K**). Statistical graphs of relative fluorescence intensities for TMRE (**L**) and MitoSOX (**M**). n = 6 (**H-I**) or n = 4 (**J**). **(N).** In ALV-J-infected cells overexpressing *HSPD1*, western blotting was performed to detect the expression levels of Parkin, TFAM, TOMM20, cytochrome *c*, and VDAC1 in mitochondrial and cytosolic fractions. n = 3. For **D-G** and **K-M**, the values are shown as the mean ± SD. *p < 0.05, **p < 0.01, ***p < 0.001 by unpaired Student’s t test.

Next, to determine whether HSPD1 has antiviral effects similar to CREG1, we independently assessed its impact on viral replication. The results of viral replication assays showed that *HSPD1* overexpression inhibited viral replication, whereas *HSPD1* knockdown significantly enhanced viral replication (Figure 6C-6D and Figure S8C-S8F). HSPD1 similarly promotes the expression of I-IFN and ISGs during the antiviral response (Figure 6E and 6F). Furthermore, under viral infection conditions, we investigated the impact of HSPD1 on mitochondrial functions. Our results demonstrate that ectopic expression of *HSPD1* indeed affects various aspects of mitochondrial biology, including a reduction in mitochondrial membrane potential (as indicated by JC-1 and TMRE) (Figure 6H-6I and 6K-6L), decreased MitoSOX fluorescence (Figure 6J and 6M), increased levels of cytosolic mtDNA (Figure 6G) and cytochrome *c* (Figure 6N), and enhanced mitophagy (Figure 7A). In contrast, knockdown of *HSPD1* led to opposite effects compared to its overexpression (Figure 6H-6K, and Figure S8G-S8J). Although ectopic expression of *HSPD1*, similar to CREG1, increased cytosolic mtDNA and cytochrome *c*, its impact on apoptosis during the antiviral response remained minimal, regardless of whether HSPD1 was overexpressed or knocked down (Figure S8K and S8L).

**Figure 7.**
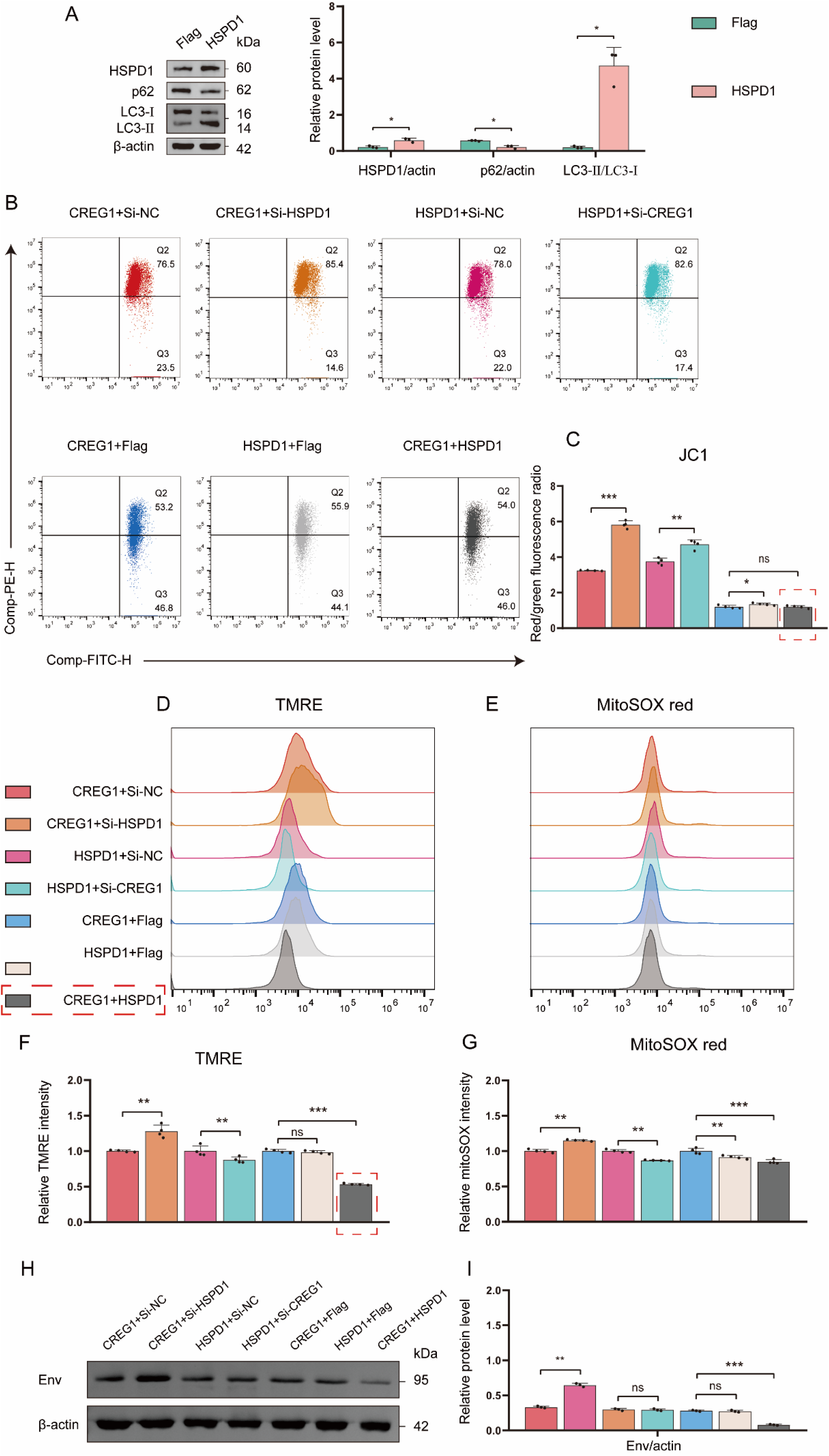
HSPD1 is required for CREG1-mediated regulation of mitochondrial function. **(A).** Western blot analysis of HSPD1, p62, and LC3 in *HSPD1*-overexpressing cells. n = 3. **(B-I).** In cells with *HSPD1* knockdown and *CREG1* overexpression, *CREG1* knockdown and *HSPD1* overexpression, or co-expression of *CREG1* and *HSPD1*, JC-1 (**B**), TMRE (**D**), and MitoSOX (**E**) fluorescence intensities were measured using a microplate reader at 48 hpi following ALV-J infection. Expression of the viral envelope protein Env was detected by western blot (**H**). The ratio of red fluorescence to green fluorescence (**C**). Statistical graphs of relative fluorescence intensities for TMRE (**F**) and MitoSOX (**G**). The relative protein quantification chart (**I**). n = 4 (**B**, **D** and **E**) or n = 3 (**I**). For **A**, **C**, **F**, **G**, and **I**, the data are presented as mean ±SD. Statistical significance was determined using unpaired Student’s t-test, with ns indicating no significant difference, *p < 0.05, **p < 0.01, and ***p < 0.001.

Finally, to determine whether the antiviral effect of CREG1 depends on HSPD1, we co-transfected Flag-CREG1 with either si-HSPD1 or HA-HSPD1, and assessed the effects of CREG1 on viral replication and mitochondrial function under conditions of *HSPD1* knockdown or overexpression. The results showed that knockdown of *HSPD1* reversed the decrease in membrane potential induced by CREG1 overexpression but did not affect CREG1-mediated antiviral activity or apoptosis. This finding suggests that the regulatory effect of CREG1 on mitochondrial function is dependent on its interaction with HSPD1 (Figure 7B-7I and Figure S9A-9C). This suggests that the regulation of mitochondrial function by CREG1 is dependent on its interaction with HSPD1. Interestingly, compared with single transfection of either gene, co-expression of *CREG1* and *HSPD1* produced a stronger antiviral effect, which was attributed to the enhanced induction of I-IFN and ISGs (Fig 7H and Figure S9C-S9H). Additionally, co-expression of *CREG1* and *HSPD1* led to a further decrease in mitochondrial membrane potential and a higher proportion of early apoptosis (Fig 7C, 7F, and Figure S9A-S9B). Collectively, these data indicate that HSPD1 is required for CREG1-mediated regulation of mitochondrial function, but not for antiviral activity and apoptosis.

## Discussion

Mitochondria orchestrate innate immune defenses against RNA viruses through dynamic regulation of antiviral signaling cascades[7]. In this study, we identified CREG1 as a novel antiviral gene. The antiviral activity of CREG1 primarily derives from its regulation of mitochondrial function. Overexpression of *CREG1* leads to a decrease in mitochondrial membrane potential, triggers mitophagy, reduces ATP production and respiratory chain activity, increases ROS generation, and promotes the release of mtDNA and cytochrome c into the cytoplasm. On one hand, the release of mtDNA into the cytoplasm activates the cGAS-STING signaling pathway, resulting in increased I-IFN production and upregulation of interferon-stimulated genes. On the other hand, the elevated intracellular levels of reactive oxygen species (ROS) and the release of cytochrome c into the cytoplasm, which induce apoptosis, also exert detrimental effects on viral replication. Importantly, the regulatory effect of CREG1 on mitochondrial function is mediated through its interaction with the mitochondrial chaperone HSPD1 (Figure 8). The co-expression of CREG1 and HSPD1 confers enhanced antiviral activity.

**Figure 8.**
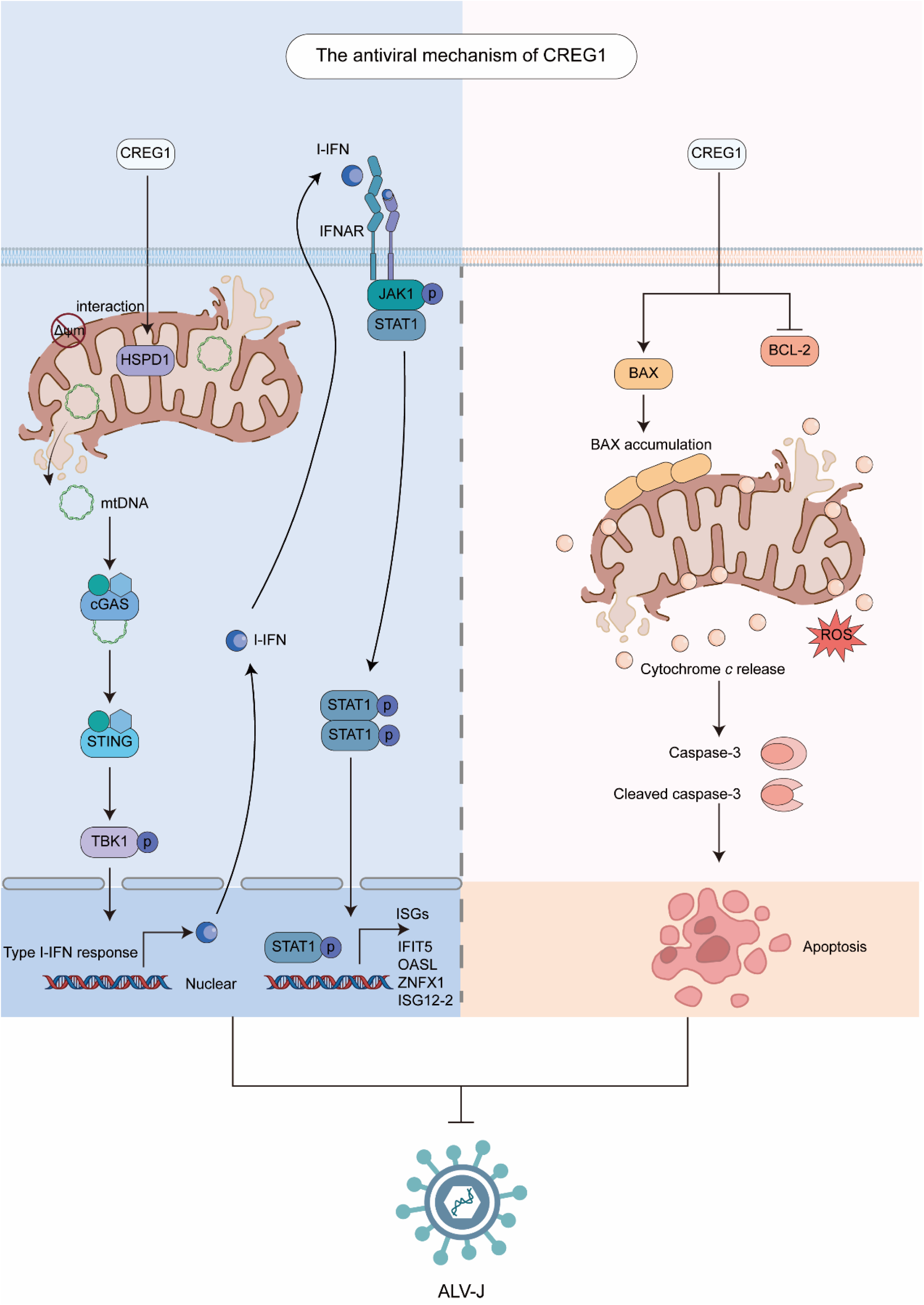
Diagram of the antiviral mechanism of CREG1.

A striking observation in our study is that CREG1-induced mitochondrial dysfunction triggers activation of the cGAS-STING signaling pathway and enhances innate immune responses via cytosolic release of mtDNA. However, we found that, compared to the control group, there was no significant increase in mtRNA release. We speculate that CREG1 induces mitochondrial membrane damage or mitochondrial stress. However, based on our TEM results, complete mitochondrial rupture did not occur. In this context, mtDNA, as a structurally stable, non-translated circular double-stranded molecule, can be more readily released even in the absence of full mitochondrial disruption[33]. In contrast, when mitochondrial damage is relatively mild, such as involving only outer membrane rupture or pore formation, large RNA molecules may not be able to pass through efficiently[34].

Meanwhile, the activation of the cGAS-STING pathway in suppressing ALV-J replication was also an unexpected finding in our study. In general, the innate immune response mediated by the cGAS-STING pathway predominantly detects double-stranded DNA (dsDNA) originating from DNA viruses, including herpes simplex virus type 1 (HSV-1) and vaccinia virus[4, 35]. However, RNA viruses can also antagonize cGAS-STING[36]. For example, in the context of chikungunya virus (CHIKV) infection, expression of the viral capsid protein alone has been shown to trigger autophagy-mediated degradation of cGAS[37]. Similarly, SARS-CoV-2 3CL protease disrupts the assembly of the functional STING signaling complex and downstream signaling by inhibiting K63-linked ubiquitination of STING[38]. In our previous studies, we speculated that MDA5 might serve as a pattern recognition receptor sensing ALV invasion[39]. However, the precise role of the cGAS-STING signaling pathway during ALV infection remains unclear. For instance, it is not known whether the lack of cGAS-STING activation is due to active antagonism by ALV viral proteins, similar to the mechanism observed in SARS-CoV-2 infection. In this study, activation of the cGAS-STING signaling pathway was found to influence ALV replication to a certain extent. These findings undoubtedly offer a novel perspective, suggesting that targeted activation of the cGAS-STING signaling pathway may serve as a potential strategy to inhibit ALV infection.

Viruses often reprogram mitochondrial respiratory activity and apoptotic signaling to create a cellular environment conducive to their replication and persistence[2]. Following SARS-CoV-2 infection, mitochondrial morphology undergoes significant alterations, characterized by matrix condensation and cristae swelling, which are associated with reduced oxidative phosphorylation (OXPHOS) polypeptides, impaired import of inner mitochondrial membrane proteins, increased mitochondrial reactive oxygen species (mROS) production, and suppression of core mitochondrial gene expression[40–43]. Adenoviral E1B 19K inhibits p53-dependent apoptosis by interacting with the pro-apoptotic protein Bak, thereby preventing the assembly of Bax and Bak on the mitochondrial membrane and suppressing mitochondria-dependent intrinsic apoptosis[44]. Similarly, vaccinia virus protein F1L has been found to inhibit apoptosis by preventing mitochondrial translocation and oligomerization of Bax on the mitochondrial membrane[45]. Cytomegalovirus (CMV) enhances cell viability during infection by utilizing the gene product of UL37, the viral mitochondrial-localized inhibitor of apoptosis (vMIA), which directly targets the cellular apoptotic mediators Bax and Bak[46–48]. However, studies on the interaction between ALV and mitochondria during infection remain limited. ALV-J is a retrovirus capable of inducing tumorigenesis in chicken tissues. We speculate that mitochondrial activity is inevitably involved in the course of viral infection, as tumorigenesis is often accompanied by mitochondrial metabolic dysfunction. Such alterations include impaired OXPHOS and elevated mROS production. The accumulation of ROS can lead to DNA damage and genomic instability, and subsequently activate oncogenic signaling pathways such as NF-κB and MAPK, thereby facilitating virus-mediated cellular transformation and tumor development[49]. Previous reports have indicated that ALV-J can activate signaling cascades such as PI3K/Akt, Wnt/β-catenin, and ERK/MAPK, which directly or indirectly regulate cell apoptosis, thereby promoting viral replication[50]. Moreover, ALV replicates slowly and establishes long-term latency within host cells, relying on the survival of infected cells to support viral replication or reactivation. Thus, the inhibition of apoptosis is essential for efficient ALV replication. For example, chicken telomerase reverse transcriptase (chTERT) can suppress apoptosis by downregulating the expression of Caspase-3, Caspase-9, and BAX, thereby promoting ALV replication in LMH cells[51]. In this study, CREG1 induced robust apoptosis by promoting mitochondrial dysfunction, which led to the release of cytochrome *c* into the cytosol. Notably, treatment with an apoptosis inhibitor partially attenuated the antiviral effect of CREG1. However, since our findings are limited to the cellular level, and ALV infection in vivo involves more complex host–virus interactions, it would be an oversimplification to conclude that promoting apoptosis alone is sufficient to inhibit viral infection. Future studies should further explore the interplay between ALV infection, mitochondrial function, and apoptosis, which may provide new strategies for combating ALV.

Whether through activation of the cGAS-STING signaling pathway or induction of apoptosis to initiate cellular immunity, the antiviral activity of CREG1 stems from its regulation of mitochondrial function. Although CREG1 has been reported to localize to mitochondria[24], our results indicate that it is primarily a cytoplasmic or nuclear protein. Its ability to modulate mitochondrial function fundamentally relies on its interaction with the mitochondrial chaperone protein HSPD1. As a mitochondrial chaperone, HSPD1 assists in the import and proper folding of proteins within mitochondria, playing a critical role in metabolic reprogramming[52]. Loss of HSPD1 triggers the mitochondrial unfolded protein response (mtUPR), resulting in impaired mitochondrial function and compromised cellular stemness[53]. Beyond its classical function in protein quality control, HSPD1 has emerged as a multifunctional molecule involved in various disease processes. For example, the baculoviral protein LEF-11 has been reported to hijack host factors ATAD3A and HSPD1, thereby facilitating efficient viral replication[54]. In numerous cancers, including pancreatic ductal adenocarcinoma[55], head and neck cancer[56], and glioblastoma[57], HSPD1 is frequently overexpressed and contributes to tumor progression. It supports OXPHOS and ATP production, thereby sustaining the high metabolic demands of rapidly proliferating tumor cells. In this study, similar to CREG1, overexpression of *HSPD1* also exhibited the ability to inhibit ALV-J replication, primarily by promoting the expression of IFN-β and ISGs. Importantly, previous studies have reported that HSPD1 interacts with IRF3 to promote the induction of IFN-β[58]. Our results indicate that *HSPD1* overexpression leads to mitochondrial dysfunction, which in turn promotes IFN-β expression. Nevertheless, whether HSPD1 interacts with members of the IRF family in this context remains to be elucidated.

Furthermore, knockdown of *HSPD1* reversed mitochondrial dysfunction induced by CREG1, clearly demonstrating that the regulation of mitochondrial function by CREG1 depends on its interaction with HSPD1. Regrettably, despite the dependence of mitochondrial regulation by CREG1 on HSPD1, the underlying mechanism explaining why the effect of CREG1 on apoptosis remains largely unaffected following *HSPD1* knockdown remains unclear. In the context of cancer, HSPD1 exerts broad anti-apoptotic functions by inhibiting p53, preventing mitochondrial permeability transition, and activating the IKK/NF-κB survival pathway[59, 60]. Nevertheless, during ALV-J infection, modulation of *HSPD1* expression, either by overexpression or knockdown, exerts only a minor impact on apoptosis. We speculate that CREG1 may regulate apoptosis by influencing other mitochondria-related proteins, as our Co-IP analysis of potential CREG1-interacting proteins identified multiple mitochondrial-associated proteins, such as SLC25A4. In a recent report, SLC25A4 was identified as a novel regulator of mitochondrial dysfunction and apoptosis-related cardiomyocyte subpopulations. Moreover, downregulation of SLC25A4 effectively improved mitochondrial function and reduced apoptosis[61]. Similarly, PHB and PHB2 have also been found to regulate apoptosis by modulating mitochondrial function[62, 63]. These observations suggest that CREG1 may trigger apoptosis through a broader mitochondrial regulatory network beyond its interaction with HSPD1.

Finally, we were surprised to find that co-expression of CREG1 and HSPD1 resulted in a more potent antiviral effect, accompanied by higher expression levels of IFN-β and ISGs. Interestingly, a previous study using a lentiviral library expressing 383 individual ISGs identified 69 ISGs that synergized with ZAP to exert antiviral effects. Further analyses demonstrated that 31 of these ISGs exhibited significant synergy with ZAP, and quantitative data showed that co-expression of these ISGs with ZAP led to a substantial reduction in viral infection rates, markedly outperforming the effect of either factor alone[64]. Given that our previous work validated the antiviral activity of ISGs such as ACSL1[65] and CH25H[66], the potential for synergistic interactions among antiviral effectors to enhance host defense warrants further investigation.

Collectively, our study uncovers CREG1 as a novel antiviral gene that may serve as a potential target for controlling ALV infection. Finally, we suggest that in-depth investigation of the crosstalk between viral infection, mitochondrial function, innate immune responses, and apoptotic pathways could provide novel strategies for the control of ALV.

## Materials and methods

### Pathological chicken tissues

The chicken tissues used for transcriptome sequencing or quantitative reverse transcription polymerase chain reaction (RT-qPCR) in this study were previously collected[25], initially preserved in liquid nitrogen, and subsequently stored at −80°C. The ALV-J-infected and control chickens had been validated in our previous study[25].

### Cell culture

DF-1 and HD11 cells, preserved in our laboratory, and chicken sarcoma bone 1 (MSB-1)cells, purchased from the American Type Culture Collection (ATCC, USA), were used in this study. All the cells were cultured at 37 °C and 5 % CO_2_. HD11 cells were cultured in Roswell Park Memorial Institute (RPMI) 1640 medium (Gibco, # 11875093). DF-1 and MSB-1 cells were cultured in Dulbecco’s modified Eagle’s medium (DMEM, Gibco, # 11965092). 10% fetal bovine serum (FBS, ABW, # AB-FBS-1050S), 5% chicken serum (only for MSB-1 cells), 100U/mL penicillin and 100µg/mL streptomycin were supplied to the base medium.

### Virus infection

The ALV-J strain, SCAU-HN06, was kindly provided by Professor Cao Weisheng from South China Agricultural University. A 10^4^ TCID_50_/0.1 mL of ALV-J SCAU-HN06 were used in this study. Cells were infected with ALV-J SCAU-HN06 in without DMEM. Following 2h of incubation, washed by 1×PBS, and the media was replaced with DMEM, supplemented with 1% FBS.

### Plasmids and RNA interference

The coding sequence (CDS) of CREG1, ITGAV or HSPD1 was cloned into the pcDNA3.1 expression vector, which contains an N-terminal ATG start codon and either a 3×Flag tag (Flag-CREG1/ITGAV) or a 3×HA tag (HA-HSPD1). Small interfering RNAs (siRNAs) targeting *ZEB2*, *BCL2L10*, *ITGAV*, *CTSB*, *CREG1*, *HSPD1*, as well as negative control RNAs (siNC), were synthesized by Gene Create Company (Wuhan, China). Cells were transfected with plasmids or siRNAs by lipofectamine 3000 (Invitrogen, # L3000015) according to the manufacturers’ instructions. The overexpression or knockdown efficiency was checked 36-48 h after transfection using RT-qPCR. Targeting sequences were listed in Supplementary Table 1.

### RNA isolation and RT-qPCR

Total RNA was isolated using TRIzol reagent (Invitrogen, # 15596026), followed by cDNA synthesis with reverse transcription kit (EZBioscience, A0010CGQ) and diluted 5-fold. cDNA amplification was performed using SYBR Green (Vazyme Biotech, # Q713-02) with the QuantStudio^TM^ 5 Real-Time PCR System (Thermo Fisher Scientific, USA). The results were calculated by 2^ΔΔCt^ method with the normalization to GAPDH. All primers were synthesized from Tsingke Biological Technology (Beijing, China) and primers sequences were shown in Supplementary Table 2.

### Western blotting

Total protein was extracted using IP lysis buffer containing 1% PMSF solution (Beyotime Biotech, # ST506) and phosphatase inhibitors (MCE, # HY-K0021). Quantification of protein was conducted with bicinchoninic acid (BCA) assay kit (Beyotime Biotech, P0009). Equal amounts of protein were separated by 10% SDS-PAGE electrophoresis (Beyotime Biotech, # P0804S) and then transferred to 0.45 μm PVDF membranes (Beyotime Biotech, # FFP22). After being blocked by 5% non-fat milk in TBST for 1h, membranes were incubated with corresponding primary antibodies at 4℃ overnight. Subsequently, the membranes were washed with TBST three times, followed by incubation with HRP-conjugated secondary antibodies for 1 h at room temperature. Signals were detected with enhanced chemiluminescence detection kit (Beyotime Biotech, # P0018M) and visualized using ODYSSEY^®^FC Imager System (LI-COR, USA) and quantified by gel analysis using ImageJ software (National Institutes of Health, USA). The antibodies used in this study were listed in Supplementary Table 3.

### RNA-seq library construction and sequencing

Total RNA was isolated using TRIzol reagent (Invitrogen). Subsequently, library construction and RNA sequencing (RNA-seq) were performed using the Illumina NovaSeq 6000 platform (Illumina, USA). Raw sequencing reads were processed to remove adaptor sequences, poly-N sequences, and low-quality reads. The cleaned reads were then mapped to GenBank to identify known chicken mRNA. RNA-seq and analysis were conducted by BIOMARKER technologies (Beijing, China).

### Immunoprecipitation (IP) and mass spectrometry

Total protein was collected from Flag-CREG1 or HA-HSPD1 overexpression cells using IP lysis buffer (Beyotime Biotech, # P0013). After centrifugation, total cellular protein was incubated with anti-Flag (Proteintech, # 66008-4-Ig) or anti-HA (Proteintech, # 51064-2-AP) M2 magnetic beads (Sigma-Aldrich, # M8823) overnight at 4°C. The precipitate was washed three times with lysis buffer and then boiled in SDS-PAGE loading buffer (Beyotime Biotech, # P0286). The supernatant was then extracted and subjected to immunoblotting with the appropriate antibodies.

Total protein was collected from Flag-CREG1 overexpression DF-1 cells using IP lysis buffer (Beyotime Biotech). The cell lysate (1000 μg) was incubated overnight at 4°C with IgG (Abcam) or Flag (Proteintech) antibodies and protein A/G beads (Thermo Fisher Scientific, # 88802). After incubation, the beads were washed and resuspended in 2× loading buffer. Each sample was separated using a 10% SDS–PAGE gel and visualized using mass spectrometry-compatible silver staining (Invitrogen). Similar conditions utilizing chicken IgG antibody (Bioss) were applied as the control lane for each gel. Mass spectrometry analyses were conducted using an LC–MS/MS system (Ekspert™ nanoLC, ABSciex Triple TOF™ 5600-plus) to identify proteins potentially interacting with CREG1.

### Data independent acquisition (DIA) quantitative proteomic techniques

Overexpression of *CREG1* or the control group EGFP in DF-1 cells, followed by ALV-J infection at 24 h, with cells harvested at 48 h post-infection. DIA quantitative proteomic techniques were performed as previously described[67]. DIA analysis was provided by Guangzhou Banianmedical Biotechnology Co., Ltd. (Guangzhou, China).

### Confocal Microscope

DF-1 cells were transfected with the Flag-CREG1 expression plasmid for 24h, then infected with ALV-J for an additional 24h. Cells were harvested and fixed in 4% paraformaldehyde, permeabilized with 0.1% saponin, blocked for 30 min with 10% goat serum, and incubated anti-Flag at 4°C overnight. After each step, it is necessary to wash the cells three times with a washing solution. Subsequently, samples were incubated FITC-labeled Goat Anti-Mouse IgG (Beyotime Biotech, # A0568) for 1h, and DAPI was used to stain the nuclei. Images were visualized by confocal laser microscopy (Leica).

### Detection of mitophagy by confocal microscopy

To evaluate mitophagy, cells were co-infected with Ad-GFP-LC3 (Hanbio, # HBLV-1010) for 24 h, followed by staining with MitoTracker Red (200 nM, Beyotime, # C1032) at 37°C for 30 min in the dark. Images were acquired using a laser scanning confocal microscope (Leica). Colocalization indicates the occurrence of mitophagy.

### Transmission electron microscopy (TEM)

Cells were transfected with Flag-CREG1, si-CREG1, or the corresponding controls for 24 h, followed by ALV-J infection for 48 h. Cells were harvested and fixed using 2% glutaraldehyde for 24 h at 4°C. The samples underwent various processing steps, including fixation, gradient alcohol dehydration, and displacement, before being imaged with a transmission electron microscope (JEM-2000EX TEM, Japan).

### Mitochondrial membrane potential, ROS, and superoxide detection

#### JC-1 staining

Mitochondrial membrane potential (Δψm) was assessed using the JC-1 dye (Beyotime Biotech, # C2003S) according to the manufacturer’s instructions. Briefly, cells were harvested and incubated with JC-1 working solution at 37°C for 20 min in the dark. After washing twice with JC-1 buffer, fluorescence was measured by flow cytometry. Mitochondrial depolarization was indicated by a decrease in the red/green fluorescence intensity ratio.

#### MitoROS detection

MitoROS was assessed using the MitoSOX™ Red mitochondrial superoxide indicator (Beyotime Biotech, #S0061S). Cells adhering to coverslips in a 35 mm dish were treated with 2 mL of the working solution of MitoSOX reagent for 30 min at 37°C and 5% CO2. After incubation, cells were washed three times with PBS. The fluorescence intensity was measured using flow cytometry with excitation/emission at 510/580 nm.

#### TMRE staining

TMRE (tetramethylrhodamine ethyl ester) staining was used to evaluate mitochondrial membrane potential. Cells were incubated with 200 nM TMRE (Beyotime Biotech, #C2001S) at 37°C for 20 min in the dark, then washed with PBS. The fluorescence intensity was detected by flow cytometry using the PE channel (excitation/emission: 550/575 nm).

### Flow Cytometry for detecting apoptosis

The prepared cells were added to 500 mL of 1× Annexin V buffer and gently resuspended. The cell suspensions were then incubated with 5 mL of Annexin V-FITC/APC (Beyotime Biotech, # C1383S)/(Abcam, # ab236215) and 5 mL of propidium iodide staining solution at 25°C for 15 min in the dark before being analyzed by flow cytometry.

### Generation of mitochondria-deficient ρ⁰ cells

Cells were treated with 50 ng/mL ethidium bromide (Sigma, # E1510), supplemented with 50 μg/mL uridine (MCE, HY-B1449) and 1 mM sodium pyruvate (MCE, HY-W015913) for 5 d, as previously described[68]. Control cells were cultured under identical conditions without exposure to ethidium bromide. The deletion of mitochondrial DNA was verified by staining cells with MitoTracker Red (Beyotime, # C1032) or by RT-qPCR to detect the total cellular mtDNA content.

### Mitochondrial cytosolic separation

The cell mitochondria isolation kit (Beyotime Biotech, # C3601) is used for isolating mitochondria and cytoplasm from cultured cells. Briefly, collect the processed cells, wash once with cold PBS, then add 1-2.5 mL of mitochondrial isolation reagent to 20-50 million cells, gently resuspend the cells, and incubate on ice for 10-15 min. Transfer the cell suspension to a suitably sized glass homogenizer and homogenize for 10-30 strokes. Carefully transfer the homogenate to a new centrifuge tube and centrifuge at 11,000 g for 10 min at 4°C. Carefully remove the supernatant. The pellet contains the isolated mitochondria.

### Cytosolic DNA extraction

Cytosolic extracts were isolated primarily as previously described[69]. Briefly, cells were resuspended in 500 µL of buffer containing 150 mM NaCl, 50 mM HEPES, and 40 µg/mL Digitonin, followed by agitation at 4 °C for 15 min. The homogenates were centrifuged at 2000×g for 5 min, and the supernatant, containing the cytosolic fraction, was transferred to a fresh tube. The pellet from this centrifugation was saved for western blot analysis. To ensure the cytosolic fraction was free of nuclear, membrane, and mitochondrial contaminants, it was further purified by three additional rounds of centrifugation at 2000×g. The purity of both the cytosolic and pellet fractions was verified by western blotting. Total DNA was extracted from our cytosolic fraction using DNA extraction kit (Omega).

### ROS assay

To determine ROS production, cells were treated with 10 mM 2’,7’-Dichlorodihydrofluorescein (DCFH-DA) following the manufacturer’s protocols (Beyotime Biotech, # S0035S). Set the microplate reader to an excitation wavelength of 488 nm and an emission wavelength of 525 nm to measure the fluorescence intensity of DCF.

### Quantification of intracellular nitric oxide (NO) levels

Cellular nitric oxide (NO) levels were determined using a commercial detection kit based on nitrate reductase activity (Beyotime, Cat# S0024) following the supplier’s guidelines. After treatment, cells were collected, lysed, and centrifuged to obtain the supernatants. Nitrate in the samples was enzymatically converted to nitrite, which then reacted with Griess reagent to generate a colored product. The absorbance at 540 nm was recorded using a microplate spectrophotometer, and NO concentrations were quantified using a sodium nitrite standard curve.

### Measurement of intracellular ATP levels

ATP levels were measured using an ATP Assay Kit (Beyotime, Cat# S0027) according to the manufacturer’s instructions. Briefly, cells were lysed with the lysis buffer provided in the kit, and the lysates were centrifuged at 12,000 × *g* for 5 min at 4 °C. The supernatants were collected, and equal volumes of the samples and the luciferase reagent were mixed in a white 96-well plate. Luminescence was immediately measured using a microplate reader. ATP concentrations were calculated based on a standard curve and normalized to the total protein content of each sample.

### Measurement of NADH-CoQ reductase and Cytochrome C Oxidase activities

The activities of NADH-CoQ reductase (Complex I) and Cytochrome C Oxidase (Complex IV) were determined using commercial assay kits from Solarbio (Cat# BC0510 and Cat# BC0940) following the manufacturer’s protocols. Briefly, cells were collected and lysed using the extraction buffer provided in the kit under ultrasonic homogenization. After centrifugation, the enzyme activity was measured using the resulting supernatant. The absorbance changes were recorded at the specified wavelength using a microplate reader. Enzyme activities were calculated based on the extinction coefficient and normalized to the total protein content.

### Enzyme-linked immunosorbent assay (ELISA)

The supernatants were collected from *CREG1*-overexpressing or *CREG1*-knockdown cells 48 h post ALV-J infection. The release of chicken IL-1β (Mlbio, # ml042758), TNF-α (Mlbio, # ml002790), IL-10 (Mlbio, # ml059830), IL-4 (Mlbio, # ml059838), IFN-γ (Mlbio, # ml023435), IFN-β (Mlbio, # ml059828), and cGAMP (Cayman chemical, # 501700) was measured using quantitative ELISA kits according to the manufacturer’s instructions.

### Statistical analysis

Statistical analysis was performed using GraphPad Prism 8 software (GraphPad Software, Inc.). All data in this study were generated from at least three independent replicated experiments. Data are presented as the mean ± standard deviation (SD). Statistical significance was tested using the two-tailed Student’s *t* test and is indicated by *p* values; *p < 0.05, **p < 0.01 and ***p < 0.001 were considered to indicate significance.

## Acknowledgement

We note our admiration and respect for researchers in this field and in our laboratories, for their dedication and hard work. This work was supported by the National Natural Science Foundation of China (grant numbers 31801030 and 31571269), the China Agriculture Research System (grant number CARS-41).

## Ethics statement

All animal research projects were sanctioned by the South China Agriculture University Institutional Animal Care and Use Committee. All animal procedures were performed according to the regulations and guidelines established by this committee and international standards for animal welfare.

## Author contributions

Q.Z., W.L, and X.Z. conceived and designed the experiments, supervised the research process, and drafted the manuscript. Q.Z., M.W (Meihuizi Wang)., M.P (Ming Pan)., J.X (Junliang Xia)., T.X. (Tao Xu) performed all experiments, conducted data analysis and statistical work. Q.Z., W.L, and X.Z. revised the article. X.Z. approved the final manuscript.

## Data availability

All the data required to support the conclusions of this paper are included within the main text and/or the Supplementary Materials. Original images of the Western blots and qPCR results are provided as supplementary content.

